# The single-cell epigenetic regulatory landscape in mammalian perinatal testis development

**DOI:** 10.1101/2021.03.17.435776

**Authors:** Jinyue Liao, Hoi Ching Suen, Shitao Rao, Alfred Chun Shui Luk, Ruoyu Zhang, Annie Wing Tung Lee, Ting Hei Thomas Chan, Man Yee Cheung, Ho Ting Chu, Hon Cheong So, Robin M. Hobbs, Tin-Lap Lee

**Affiliations:** Developmental and Regenerative Biology Program, School of Biomedical Sciences, Faculty of Medicine, The Chinese University of Hong Kong, Shatin, Hong Kong SAR, China; Cancer Biology and Experimental Therapeutics Program, School of Biomedical Sciences, Faculty of Medicine, The Chinese University of Hong Kong, Shatin, Hong Kong SAR, China; School of Medical Technology and Engineering, Fujian Medical University, Fujian, P.R. China; Germline Stem Cell Biology Laboratory, Centre for Reproductive Health, Hudson Institute of Medical Research, Australia

## Abstract

Spermatogenesis depends on an orchestrated series of developing events in germ cells and full maturation of the somatic microenvironment. To date, the majority of efforts to study cellular heterogeneity in testis has been focused on single-cell gene expression rather than the chromatin landscape shaping gene expression. To advance our understanding of the regulatory programs underlying testicular cell types, we analyzed single-cell chromatin accessibility profiles in more than 25,000 cells from mouse developing testis. We showed that scATAC-Seq allowed us to deconvolve distinct cell populations and identify cis-regulatory elements (CREs) underlying cell type specification. We identified sets of transcription factors associated with cell type-specific accessibility, revealing novel regulators of cell fate specification and maintenance. Pseudotime reconstruction revealed detailed regulatory dynamics coordinating the sequential developmental progressions of germ cells and somatic cells. This high-resolution data also revealed putative stem cells within the Sertoli and Leydig cell populations. Further, we defined candidate target cell types and genes of several GWAS signals, including those associated with testosterone levels and coronary artery disease. Collectively, our data provide a blueprint of the ‘regulon’ of the mouse male germline and supporting somatic cells.

## Introduction

Mammalian testis consists of germ cells and distinct somatic cell types that coordinately underpin the maintenance of spermatogenesis and fertility. These testicular cells display extensive developmental dynamics during the perinatal period. Primordial germ cells (PGCs) give rise to M-prospermatogonia at about embryonic day (E) 12, which enter G0 mitotic arrest at about E14 to form T1-prospermatogonia (T1-ProSG) (McCarrey 2013). Shortly after birth, T1-ProSG resume mitotic activity and begin migrating from the center of the testis cords to the basal lamina of testicular cords, and become T2-prospermatogonia (T2-ProSG). Once resident at the basement membrane, T2-ProSG generate spermatogonia including self-renewing spermatogonial stem cells (SSCs) or directly transition into differentiating spermatogonia that participate in the first round of spermatogenesis (Kluin and de Rooij 1981; Manku and Culty 2015).

Specialized somatic cells play a pivotal role in maintaining normal germ cell development and spermatogenesis. SSCs and their initial progenies reside in a niche on the basement membrane and are surrounded by Sertoli cells, which nourish SSCs. In mice, Sertoli cells actively proliferate during the neonatal period for two weeks (Vergouwen et al. 1991). Outside the seminiferous tubules, the interstitial compartment of testis mainly contains stroma, peritubular myoid cells (PTMs), Leydig cells, macrophages and vascular cells. PTMs are smooth muscle cells that distribute over the peripheral surface of the basement membrane (Maekawa et al. 1996). They are mainly involved in tubule contractions to facilitate the movement of sperm to the epididymis and secrete extracellular matrix materials (Chen et al. 2014). Leydig cells are responsible for steroidogenesis and provide critical support for spermatogenesis. In mammals, there are two types of Leydig cells, fetal Leydig cells (FLCs) and adult Leydig cells (ALCs), which develop sequentially in the testis (Shima 2019). During the perinatal period, both FLCs and stem Leydig cells (SLCs) reside in the testis. While FLCs start to degenerate after birth, they are replaced by SLCs, which are the progenitors of ALCs (Su et al. 2018).

One of the goals of developmental biology is to identify transcriptional networks that regulate cell differentiation. Recent advances in single-cell transcriptomic methods have enabled an unbiased identification of cell types and gene expression networks in testis (Green et al. 2018; Tan et al. 2020). A key remaining question is how distinct testis cell types are developed during the perinatal period. Since all cells share the same genetic information, lineage specification must be regulated by differential chromatin accessibility in a cell type-specific and dynamic manner. Thus, the understanding of the testis development, especially the cell type-specific transcription factors (TFs) is of paramount importance. scRNA-Seq provides limited information of TFs, which are usually lowly expressed. Although sequencing methods such as ATAC-Seq and DNase-Seq have been developed for profiling chromatin accessibility landscapes across samples and classification of regulatory elements in the genome, the nature of bulk measurements masks the cellular and regulatory heterogeneity in subpopulations within a given cell type (Shema et al. 2019). Moreover, uncovering regulatory elements in testicular cell types has been particularly challenging as samples are limited and heterogeneous. Notably, only one previous study examined the genome-wide chromatin accessibility in Sertoli cells, using the *Sox9* transgenic line followed by fluorescence-activated cell sorting (FACS) and DNase-Seq (Maatouk et al. 2017).

By profiling the genome-wide regulatory landscapes at a single-cell level, recent single-cell sequencing assay for transposase-accessible chromatin (scATAC-Seq) studies have demonstrated the potential to discover complex cell populations, link regulatory elements to their target genes, and map regulatory dynamics during complex cellular differentiation processes. We reasoned that the advent of new single-cell chromatin accessibility sequencing methods, combined with single-cell transcriptomic data, would be instrumental in advancing our understanding of gene regulatory networks in mammalian testis development. Here, we applied scATAC-Seq to deconvolve cell populations and identify cell type-specific epigenetic regulatory circuits during perinatal testis development. The dataset led to identification of key cell type-specific TFs, defined the cellular differentiation trajectory, and characterized regulatory dynamics of distinct cell types. Furthermore, our results shed light on the identification of target cell types for genetic variants. To enable public access to our data, we constructed the mouse testis epigenetic regulatory atlas website at http://testisatlas.s3-website-us-west-2.amazonaws.com/.

## Results

### Single-cell ATAC-Seq captures developmental and cell type-specific heterogeneity in the testis

To delineate the dynamic changes on cellular populations in a developing testis, we profiled the chromatin accessibility landscapes of mouse perinatal testis across E18.5 and postnatal stages (P0.5, P2.5 and P5.5) by scATAC-Seq (Figure 1A). These time points were chosen to represent the diversity of cell type compositions involved in the key developmental events in the testis (Figure 1B). Altogether, we profiled chromatin accessibility in 25,613 individual cells after stringent quality control filtration and heterotypic doublet removal (Supplementary Figure S1). These samples showed no clustering based on covariates such as transcription start site (TSS) enrichment and fragment size (Supplementary Figure S2A).

**Figure 1.**
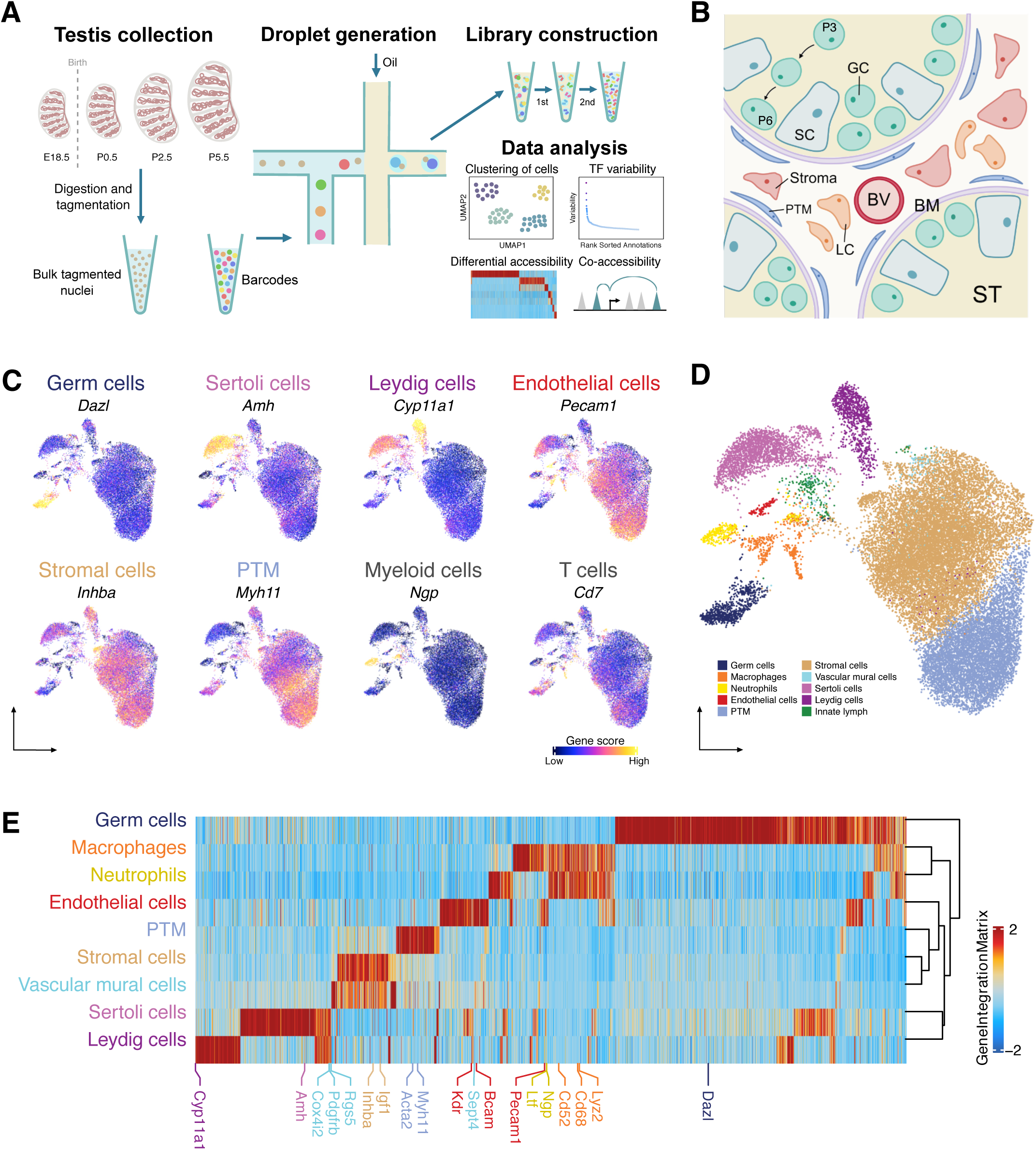
Classification and identification of germ cells and somatic cells during perinatal testicular development. A. Experimental design. The workflow of testis collection and scATAC-seq to measure single nuclei accessibility on BioRad SureCell ATAC-Seq platform. B. Illustration of the testicular microenvironment. GC: germ cell; SC: Sertoli cell; LC: Leydig cell; BV: Blood vessel; BM: Basement membrane; ST: Seminiferous tubule; PTM: Peritubular myoid cell. C. UMAP representations with cells colored by the gene score of marker genes for each cell type. D. UMAP representation of cells captured from 4 time points. Cells are colored by predicted groups. E. Heatmap of 12,250 marker genes across cell types (FDR <= 0.05, Log2FC >= 0.2).

Several clusters showed developmental stage specificity, which were made up almost entirely of cells from a single time point (Supplementary Figure S2B). To improve cell type annotation, we used Harmony to integrate datasets of 4 time points and project cells onto a shared embedding in which cells were grouped by cell type rather than developmental stage (Korsunsky et al. 2019). Unbiased iterative clustering of these single cells after integration identified 11 distinct clusters with different distribution patterns across time points (Supplementary Figure S2C and D). Some clusters could be assigned to known testicular cell types based on gene activity scores of key marker genes compiled from chromatin accessibility signals within the gene body and promoter (Tan et al. 2020) (Figure 1C).

While this approach provided broad classifications for cell type annotation, an unbiased method is needed for more accurate classification. Therefore, we leveraged a previously published scRNA-Seq dataset of perinatal testis samples to predict cell types in scATAC-Seq data (Tan et al. 2020). We first re-analyzed scRNA-Seq data to determine the cellular composition and annotate cells based on their transcriptional profiles. Prediction of cell types in scATAC-Seq was then performed by directly aligning cells from scATAC-Seq with cells from scRNA-Seq through comparing the ‘query’ gene activity scores matrix with ‘reference’ scRNA-Seq gene expression matrix. The results showed that the vast majority of cells had a high prediction score and were confidently assigned to a single cell type (Figure 1D, Supplementary Figure S2D). We further validated the cluster assignment by gene score and chromatin accessibility profiles of marker genes (Figure 1E). Taken together, scATAC-Seq allowed the detection and assignment of cell identities in the developing testis.

### Chromatin accessibility defines cell types in developing testis

Cell types can be distinguished based on whether differentially accessible chromatin regions (DARs) are ‘open’ or ‘closed’. After identifying 214,890 accessible chromatin regions in the scATAC-Seq library (Supplementary Table S1), we investigated cell type-specific chromatin accessibility profiles. We compared differences in chromatin accessibility among cell types directly using Wilcoxon testing to identify DARs while accounting for TSS enrichment and the number of unique fragments per cell (Figure 2A and Supplementary Table S2 and 3).

**Figure 2.**
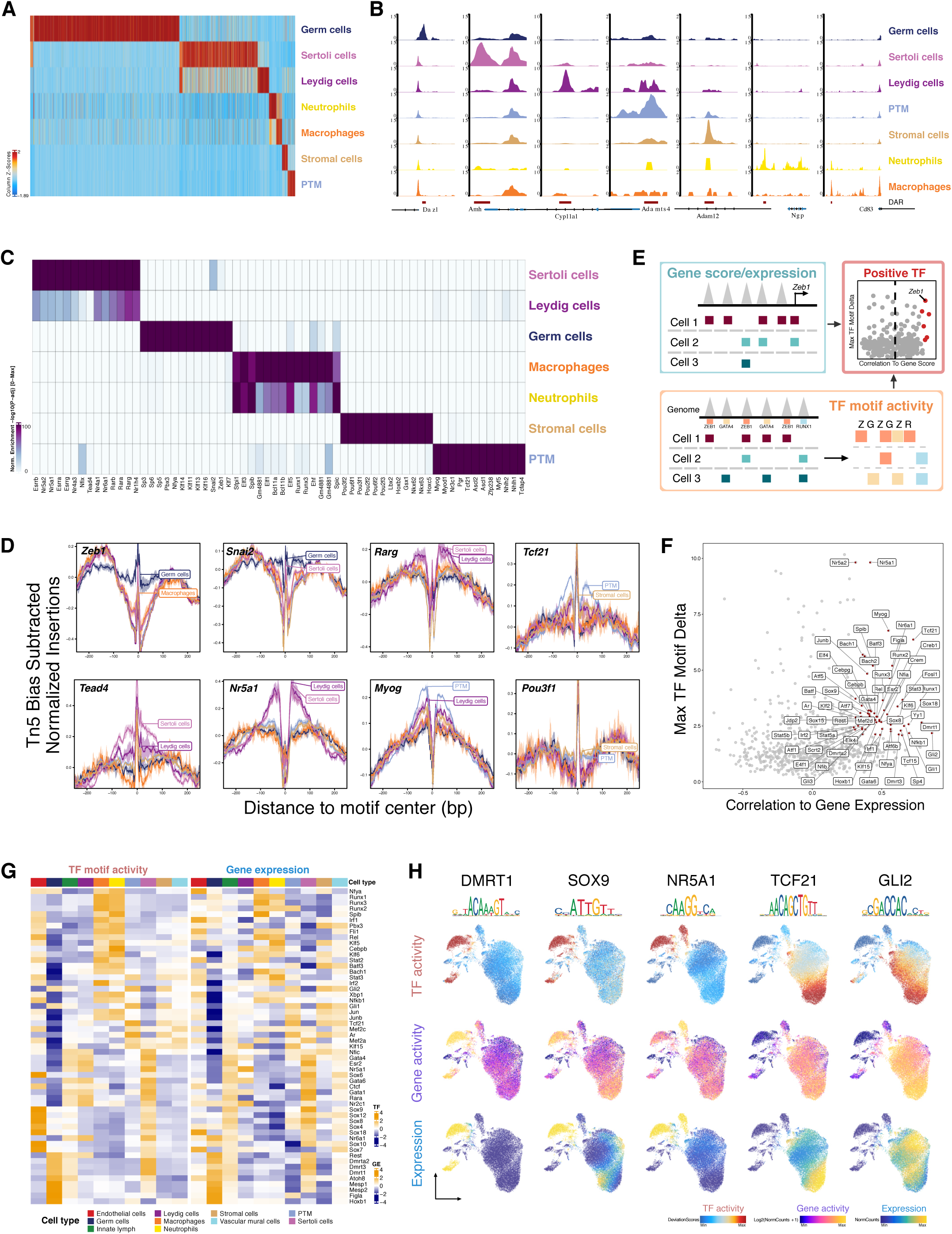
Characterization of differentially accessible regions and identification of cell type-specific transcription factors (TFs). A. Heatmap of 51,937 differentially upregulated accessible peaks (FDR <= 0.01, Log2FC >= 2) across cell types. B. Aggregated scATAC-seq profiles of selected markers. C. Heatmap of enriched motifs (FDR <= 0.1, Log2FC >= 0.5) across cell types. D. TF footprints (average ATAC-seq signal around predicted binding sites) for selected TFs. E. Schematic of identifying positive TF regulators through correlating gene score (scATAC-seq data)/gene expression (integrating scATAC-seq and scRNA-seq data) with TF motif activity (scATAC-seq data). F. Scatter plot of positive TF regulators (correlation > 0.5, adjusted p-value < 0.01). G. Heatmaps of differential TF motif activity (left) and gene expression (right) of positive TF regulators in F. H. TF overlay on scATAC UMAP of TF chromVAR deviations (top), gene activity scores (middle) and gene expression (bottom).

Deconvolution of chromatin accessibility by cell types revealed accessible sites are primarily located in the distal and intron region (>3-kb from TSS), suggesting an enrichment of gene regulatory elements (Supplementary Figure S3A).

We found cluster-specific DARs were associated with cell type-specific marker genes identified from scRNA-Seq (Figure 2B, Supplementary Figure S3B). For example, *Amh* is a marker gene in Sertoli cells, and it showed increases in both number and amplitude of ATAC peaks within its promoter and gene body. We further compared DARs to a previously-published DNase-Seq experiments in bulk Sertoli cells and found that DNase I hypersensitive sites were clearly enriched in the Sertoli cell population in our scATAC-Seq (Supplementary Figure S3C) (Maatouk et al. 2017). These data confirm that scATAC-Seq is a robust method for the detection of cell type-specific chromatin accessibility.

### Chromatin accessibility is associated with cell type-specific transcription factor activity

Currently, the identities of cell type-specific TFs involved in testis development are poorly defined. Accessibility at regulatory sites is driven by TF binding and histone modifications of local chromatin (Cui et al. 2013). To characterize the determinants of chromatin accessibility variation among cell types, we predicted TF ‘activity’ for individual cell types based on the presence of binding motifs within DARs. Assessment of enriched TFs and their cognate motifs identified several known cell type-specific regulators – including the nuclear receptors (NR4A1 and NR5A1) in Sertoli cells and Leydig cells, MYOG in PTMs, and previously uncharacterized TFs as potential cell type-specific regulators (Figure 2C). For example, we found that ZEB1 and SNAI2 motifs were enriched in germ cells, indicating they may undergo EMT-related processes in perinatal development (Liao et al. 2020). DNA bound by TFs is protected from transposition by Tn5, which can be visualized by plotting the ‘footprint’ pattern of each TF as the local chromatin accessibility surrounding the motif midpoint. Examining the footprint validated the cell type-dependant differential footprint occupancy of identified TFs (Figure 2D).

Although motif enrichment for DAR can be informative, this measurement is not calculated on a per-cell basis and they do not take into account the insertion sequence bias of Tn5. Therefore, in the second analysis approach, we used chromVAR to infer TF motif activity, which can reflect the enrichment level of the TF motif in accessible regulatory elements in a scATAC-Seq dataset. We first identified deviant TF motifs by stratifying motifs based on the degree of variation observed across clusters. Since TFs from the same family often share a similar motif, this makes it challenging to identify the specific TFs that actually drive the observed changes in chromatin accessibility. To reduce false discovery, we focused on putative positive regulators determined from the correlation between the gene expression based on scRNA-Seq dataset (or inferred gene activity based on scATAC-Seq dataset) and the chromVAR motif activity score, reasoning that expression of high-confidence TFs is correlated with their motif accessibility (Figure 2E).

Clustering analysis of positive regulators showed that diverse combinatorial TF motif landscapes were apparent across cell types and closely mirrored gene accessibility profiles of respective TFs (Figure 2F to H). There was an increased GATA1 TF ‘activity’ (motif activity) in the Sertoli cell cluster, in addition to increased chromatin accessibility in *Gata1* (gene activity) and increased *Gata1* transcription (gene expression) Figure 2G). It has been shown that mutation of GATA1 causes human cryptorchidism (Nichols et al. 2000) and its expression in Sertoli cells is conserved between human and mouse (Yomogida et al. 1994). A similar pattern was seen for DMRT1 in germ cells. DMRT1 is one of the top motifs enriched in human SSC-specific ATAC-Seq peaks (Guo et al. 2017) and loss of DMRT1 causes spermatogonia to aberrantly enter meiosis (Matson et al. 2010). chromVAR detected an enrichment of DMRT1 binding motif in the germ cell cluster, together with increased *Dmrt1* chromatin accessibility and transcription (Figure 2G). This analysis also revealed shared and unique regulatory programs across cell types. For example, PTM and stromal cells shared similar regulators, but PTM demonstrated higher activity of AR and TCF21 (Figure 2G). Similarly, NR5A1 and GATA4 were more active in both Leydig cell and Sertoli cell populations. However, only Sertoli cells showed increased activity in the SOX family.

In order to characterize the putative downstream target genes of TFs, we examined TF regulon activity with the scRNA-Seq data using SCENIC. The regulon of a TF, which contained potential direct targets of the TF, was inferred by TF-centered gene co-expression network followed by motif-based filtration. SCENIC results revealed clear cell type enrichment of regulons. Further stratification of the TFs identified both in scATAC-Seq and SCENIC revealed several highly confident regulators in each cell type (Supplementary Figure S4). For example, the regulon activities of *Sox9* and *Gata4* were enriched in Sertoli and Leydig cells respectively. SCENIC also revealed the target genes of individual TFs. The full list of regulons and their respective target genes can be found in Supplementary Table S4. Therefore, we demonstrated that integrated analysis of scATAC-Seq and scRNA-Seq datasets yielded significant insight into TF-regulatory networks of cells in the developing testis.

Importantly, we observed that individual cell types can be defined by TF ‘activity’, suggesting that cell type-specific TFs likely regulate chromatin accessibility. Collectively, these results are indicative of robust inference of TF activity at the level of single cells and reveal TF dynamics central to *cis*-regulatory specification of diverse cell states.

### Chromatin accessibility is associated with cell type-specific chromatin interaction networks

We next focused on the regions of the genome driving cell lineage-specific gene expression. As enhancers play a critical role in establishing tissue-specific gene expression patterns during development, we predicted that active enhancers would be enriched around lineage-specific genes. To test this, we used an analytical framework to link distal peaks to genes in cis, based on the coordination of chromatin accessibility and gene expression levels across cells (Figure 3A). We identified 35,245 peak-to-gene links by correlating accessibility changes of ATAC peaks within 250 kb of the gene promoter with the mRNA expression of the gene from scRNA-Seq.

**Figure 3.**
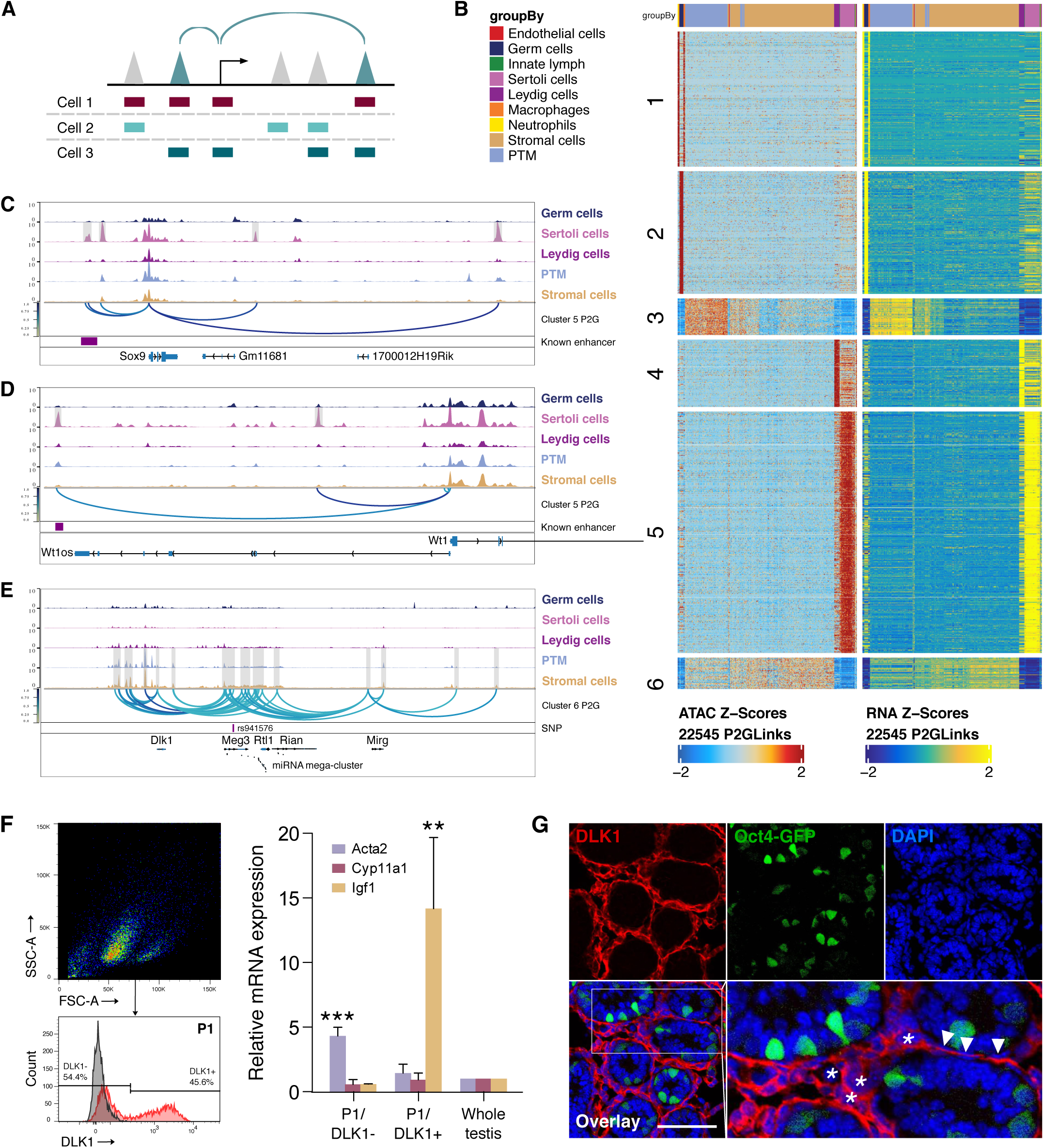
Chromatin interaction networks in different cell types. A. Schematic for identifying significant peak-to-gene links by correlating accessible peaks (scATAC-seq data) to gene expression (integrating scATAC seq data and scRNA-seq data). B. Heatmaps of TF motif activity (left) and gene expression (right) of 22,545 peak-to-gene linkages across cell types. C. Aggregated scATAC-seq profiles showing peak-to-gene links to the *Sox9* locus overlapped with known enhancer regions. D. Aggregated scATAC-seq profiles showing peak-to-gene links to the *Wt1* locus overlapped with known enhancer regions. E. Aggregated scATAC-seq profiles showing peak-to-gene links to the *Sox9* locus overlapped with SNP. F. Sorting strategy for isolation of DLK1- and DLK1+ cells from P6 whole testis. The majority of DLK1+ cells are located in P1 (upper left). The DLK1-/+ population was gated using Red-X-labelled sample compared with unstained control (lower left). RT-PCR analysis (right) of relative expression of PTM marker (*Acta2*), Leydig cell marker (*Cyp11a1*) and stromal cell marker (*Igf1*) of DLK1-/+ cells compared with whole testis sample (p < 0.001, n ≥ 3, One-way ANOVA). *Gapdh* was used as endogenous control. Error bars are plotted with SD. G. Representative confocal images of testis sections from Oct4-GFP transgenic mice at P6. Stromal cells (asterisks) and some PTM cells (arrowheads) are positive for DLK1 (red). Oct4-GFP indicated germ cells. Cell nuclei were stained with DAPI. Scale bar = 50 μm.

The information on enhancers driving gene expression in testicular somatic cell types is largely unknown. We reasoned some of the peak-to-gene links are potentially promoter-enhancer regulatory units as 3,262 peak-to-gene links were within the 17,022 annotated testis enhancers (Supplementary Figure S5A) (Gao and Qian 2020). To identify putative peak-to-gene links specific to each cell type, we performed clustering analysis, with each cluster enriched in one or more specific cell types (Figure 3B). We first performed GO analysis of the targets of peak-to-gene links and confirmed that they were highly enriched in terms related to regulations of each cell type (Supplementary Figure S5B). The full list of peak-to-gene links in each cluster can be found in Supplementary Table S5.

We next examined whether we can use this information to link DARs to known cell type-specific enhancers. During male sex determination, *Sry* activates male-specific transcription of *Sox9* in the male genital ridge via the testis-specific enhancer core element (TESCO) enhancer (Sekido and Lovell-Badge 2008). Comparison of the genomic region around *Sox9* among all cell types revealed a region 13 kb upstream formed a peak-to-gene link with the *Sox9* TSS, which is unique to the Sertoli cell population and overlapped with the 3 kb TES enhancer including the TESCO elements (Figure 3C). Enhancer activity was previously narrowed to a subregion of TES: the 1.3 kb TESCO element (Sekido and Lovell-Badge 2008). Besides known enhancers, our data linked an additional three elements located 9 kb 5′, 21 kb 3’ and 68 kb 3’ to *Sox9* representing novel regulatory elements of *Sox9* gene regulation. Our analysis also successfully revealed a previously identified functional enhancer as a novel candidate to regulate Sertoli cell marker *Wt1* (Figure 3D). After confirming that our approach can be used to reveal functionally relevant regulatory regions, we next identified putative regulatory elements for important cell type-specific regulators in each cell type, including *Nanos2* and *Uchl1* in germ cells, *Cyp11a1* and *Nr5a1* in Leydig cells, *Tpm1* and *Socs3* in PTM cells (Supplementary Figure S6), and *Dlk1* in stromal and PTM cells (Figure 3E).

Notably, the *Dlk1*-*Gtl2* locus demonstrated preferential accessibility in stromal and PTM cells. This coincided with the high number of peak-to-gene links including the largest mammalian miRNA mega-cluster located approximately 150 kb downstream (Seitz et al. 2004). We then validated the expression of *Dlk1* in mouse testicular cells. Although DLK1 is considered as a marker for immature Leydig cells in human (Lottrup et al. 2014), DLK1+ mouse testicular cells showed higher *Igf1* mRNA expression when compared with whole testis samples, indicating enrichment of stromal cells (Figure 3F). Immunostaining of neonatal testis tissue demonstrated that stromal cells and PTM cells were positive for DLK1 (Figure 3G).

In conclusion, our results highlight the occurrence of diverse cell type-specific regulatory configurations among cis-regulatory elements (CREs) and their target genes in the testis.

### Stage-specific TF regulators and chromatin co-accessibility during gonocyte to spermatogonia transition

Next, we analyzed the chromatin accessibility characteristics of the germ cells in our datasets. Re-clustering of germ cells from the embryonic day (E) 18.5, postnatal day (P) 0.5, P2.5 and P5.5 testicular datasets revealed 7 cell clusters (Figure 4A). Notably, germ cells from E18.5 and P0.5 are largely clustered together and occupy clusters C1 and C3, indicating a minimal change of chromatin accessibility before and after birth. In contrast, P2.5 cells are present in C2 and P5.5 cells occupy the remaining clusters.

**Figure 4.**
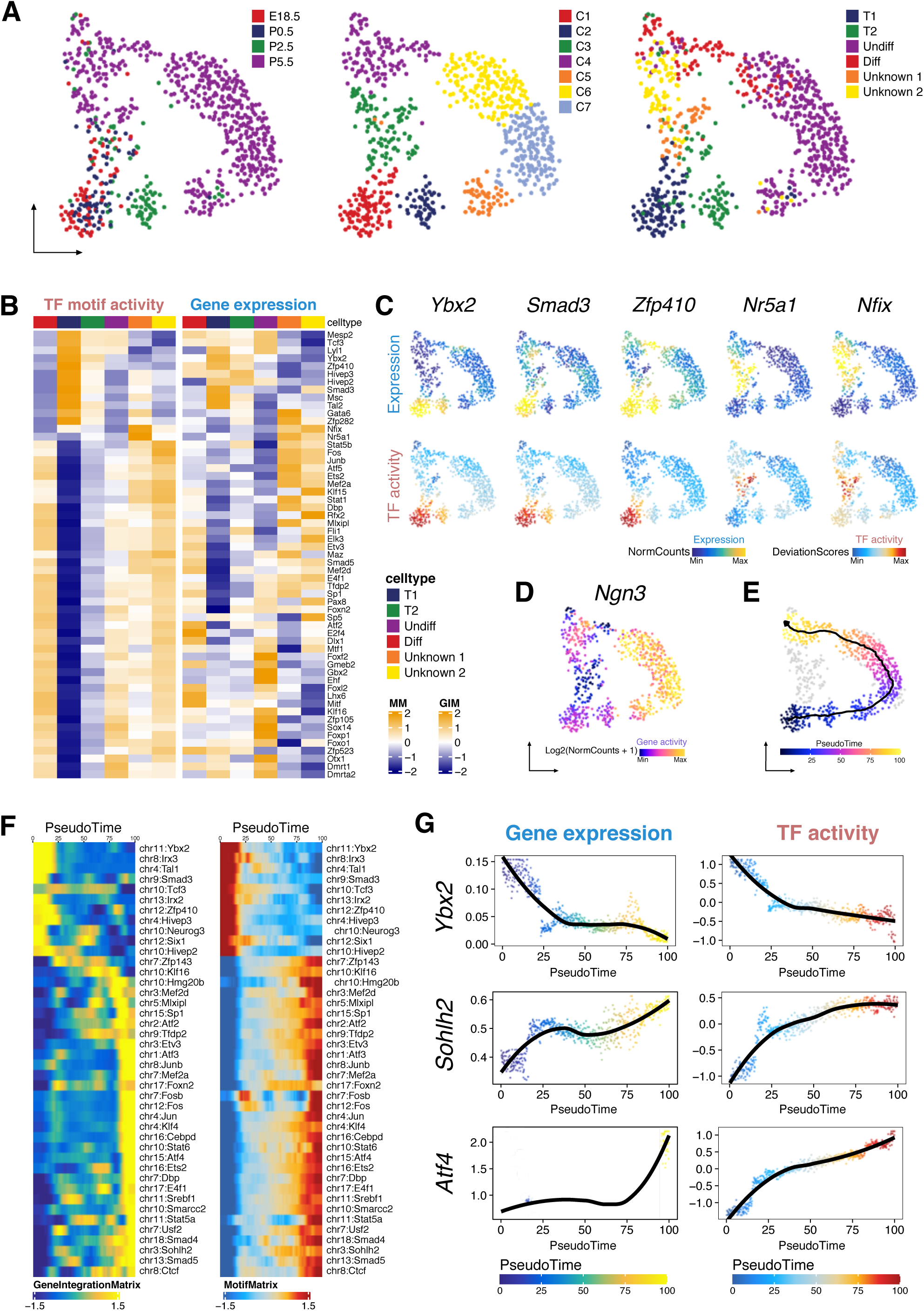
Identification of germ cell clusters during the perinatal period. A. UMAP representation of germ cells. Cells are colored by time points (left), cell clusters (middle) and clustering based on constrained integration with scRNA-seq data (right). B. Heatmaps of differential TF motif activity (left) and gene activity (right) of positive TF regulators across cell clusters (correlation > 0.5, adjusted p-value < 0.01). C. TF overlay on scATAC UMAP of gene expression (top) and TF chromVAR deviations (bottom) for positive TF regulator examples in B. D. Gene activity of *Ngn3* shown in UMAP. E. scATAC-Seq profiles are ordered by pseudotime, corresponding to the perinatal development trajectory. F. Smoothened heatmaps showing dynamic gene expression (left) and motif accessibility (right) of indicated TFs along pseudotime for gene-motif pairs of the trajectory in E. G. Gene expression (left) and motif accessibility (right) of selected TFs ordered by pseudotime.

Deconvoluting cell states using scATAC-Seq measurements alone is difficult within a single cell type. Therefore, we integrated the germ cell subsets based on scATAC-Seq data with the published perinatal testis scRNA-Seq dataset (Supplementary Figure S7A). The prediction scores of individual cells were overall high, indicating the cluster identity assignment was reliable (Supplementary Figure S7B and C). This prediction revealed 4 clusters of known developmental stages, T1-ProSG (T1), T2-ProSG (T2), undifferentiated spermatogonia (Undiff) and differentiation-primed spermatogonia (Diff), together with two clusters with unknown identity (Figure 4A).

We first identified the TFs important for each cluster, revealing 57 putative positive TF regulators in germ cell development (Figure 4B and C). For instance, *Foxo1* and *Dmrt1* exhibited increased TF motif activity and gene expression in undifferentiated spermatogonia. *Foxo1* is involved in SSC maintenance and initiating spermatogenesis (Goertz et al. 2011). *Dmrt1* prevents spermatogonia from undergoing meiosis by repressing *Stra8* transcription (Matson et al. 2010). *E2f4* shows highest activity in differentiation-primed spermatogonia, and is known to be critical for the development of the male reproductive system (Danielian et al. 2016). Our results also suggested some TF candidates regulating T1-ProSG, which have previously been difficult to identify due to technical challenges in isolating this cell population. Concordant with its role in testicular development, *Gata6* is upregulated in T1- and T2-ProSG (Padua et al. 2015). *Ybx2*, *Smad3* and *Msc* are also enriched in T1-ProSG. Lastly, we observed *Nfix* and *Nr5a1* showed increased activity in the unknown clusters.

We then aimed to reconstruct the differentiation trajectories by ordering the clusters with developmental stages predicted by scRNA-Seq integration. Two possible trajectories were observed, as germ cell differentiation appeared to diverge at P0.5 via two distinct branches. The first trajectory represents the differentiation fit into the conventional model as it charts a trajectory from gonocyte to undifferentiated and then differentiating spermatogonia in P5.5. The second path bypassed the undifferentiated state but passed through the unknown populations and directly reached the differentiating state by P5.5. It has been reported that the first round of spermatogonia arise from a unique neurogenin-3 (*Ngn3*) negative pool of ProSG that transition directly into A1 spermatogonia (Yoshida et al. 2006). Interestingly, the unknown population (Unknown-2) displayed the lowest level of *Ngn3* gene activity, which raised the possibility that the second trajectory represents the origin of the first wave of spermatogenesis (Figure 4D).

To determine the key genes driving the spermatogonial development in the first trajectory, we generated a pseudotime trajectory and uncovered a list of genes with dynamic changes (Figure 4E and Supplementary Figure S7D). The pattern of TF dynamics suggested a model of differentiation as a transition between two phases involving progressive loss of gonocyte-specific TF activities and gradual increase of TF activities relevant to spermatogonia. We observed that the motif binding activity of *Id4* was initially high but then declined after birth, while that of ETS and Sp/KLF family members increased in spermatogonia (Supplementary Figure S7D). To further reveal TFs that drive the germ cell development, we pruned the data by correlating gene expression of a TF to its corresponding TF z-score (Figure 4F and G). This method accurately identified recognized regulators in spermatogonial differentiation. For example, *Sohlh2*, which is critical for early spermatogenesis, is more accessible at the late stage (Hao et al. 2008). This also raised some novel candidates regulating spermatogonial differentiation, such as *Ybx2* and *Atf4*, which are both essential to male fertility (Fischer et al. 2004; Yang et al. 2005).

We further predicted regulatory interactions from scATAC-Seq data, identifying 12,819 putative peak-to-gene links (Supplementary Figure S8). For example, Cluster 1 included peaks linked to stem cell-related genes *Sdc4* and *Cdh1* (Supplementary Figure S9). Cluster 2 included peaks connected to genes related to progenitor and differentiating cells such as proliferation marker *Top2a*. Cluster 3 peak-to-gene links were more accessible predominantly in T1-ProSG, such as *T* and *Fbxo4*.

Taken together, our results could lead to a deeper understanding of the expression pattern of TF regulators in cells of the developing testis and also reveal candidate CREs essential for regulating spermatogonial development and differentiation.

### TF dynamics during perinatal Sertoli cell development

Re-clustering of Sertoli cells revealed 6 cell clusters (Figure 5A). Notably, C1 is largely distinct from other clusters and comprises mainly E18.5, P0.5 and P2.5 cells, suggesting this subpopulation is only present during the embryonic and early neonatal period. Based on the inferred gene scores calculated by ArchR, 81 marker genes were identified among the clusters (FDR < 0.1, log_2_FC > 0.5) (Supplementary Figure S10A). C1 is the most variated from the other clusters, with marker genes related to spermatogenesis, such as *Fzr1*, *Egfr* and *Npas2*. Clusters C4, C6, C5 and C3 consisted of cells from E18.5, P0.5, P2.5 and P5.5, respectively, which might represent sequential stages along the Sertoli cell developmental continuum. We first reconstructed the trajectory by ordering these cell clusters of the 4 time points and generated a pseudotime trajectory to uncover the critical TFs (Figure 5B). We then correlated the gene score of a TF to its corresponding TF z-score to reveal TFs enriched at different developmental stages (Figure 5C). For instance, GATA2 and PPARα were enriched at early developmental stages (Supplementary Figure S10B). GATA2 has been identified as a target of androgen receptor (Ar) in Sertoli cells, while PPARα regulates cholesterol metabolism and lipid oxidation in Sertoli cells (Bhardwaj et al. 2008; Shi et al. 2018). In contrast, HIC1 and CEBPD were upregulated at the later stage (Supplementary Figure S10B). Conditional knockout of *Hic1* in mice resulted in fewer Sertoli cells in seminiferous tubules (Uchida et al. 2020). Further, induction of C/EBP proteins by cAMP may play a role in FSH-dependent regulation in Sertoli cells (Grønning et al. 1999).

**Figure 5.**
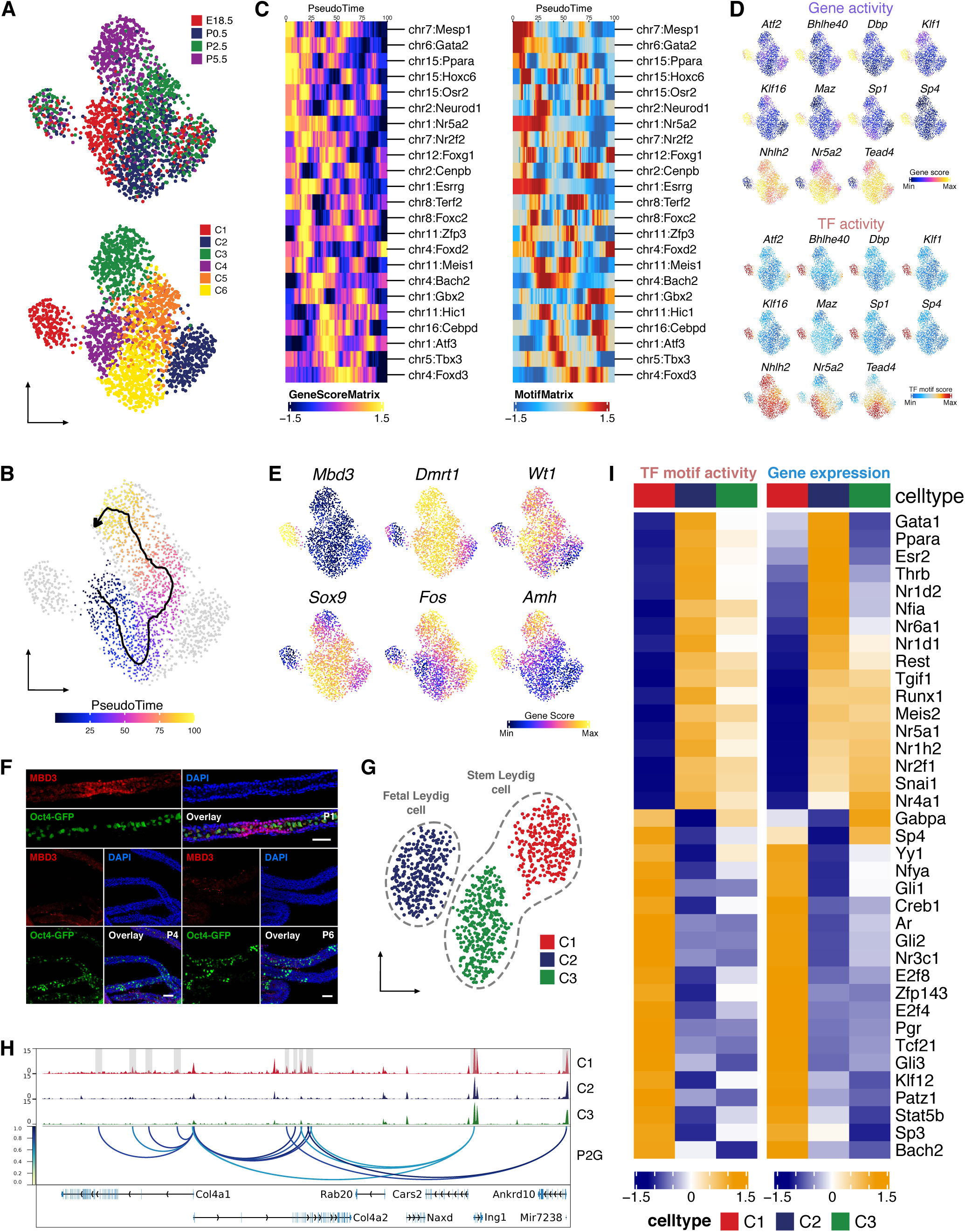
Identification of Sertoli and Leydig cell clusters during the perinatal period. A. UMAP representation of Sertoli cells. Cells are colored by time points (left) and cell clusters (right). B. scATAC-Seq profiles are ordered by pseudotime, corresponding to the perinatal development trajectory. C. Smoothened heatmaps showing dynamic gene score (left) and motif accessibility (right) of indicated TFs along pseudotime for gene-motif pairs of the trajectory in B. D. TF overlay on scATAC UMAP of gene activity scores (top) and TF chromVAR deviations (bottom) for positive TF regulators in E. E. Gene activity scores of *Mbd3*, *Fos* and Sertoli cell marker genes (*Dmrt1*, *Wt1*, *Amh* and *Sox9*) shown in UMAP. F. Representative confocal images of seminiferous tubules from Oct4-GFP transgenic mice at P1, 4 and 6. A subpopulation of Sertoli cells showed high MBD3 (red) in P1 tubules. Oct4-GFP indicated germ cells. Cell nuclei were stained with DAPI. Scale bar = 50 μm. G. UMAP representation of Leydig cells. Cells are colored by cell clusters. H. Aggregated scATAC-seq profiles showing peak-to-gene links to the *Col4a1* and *Col4a2* loci in C1 cluster. I. Heatmaps of differential TF motif activity (left) and gene expression (right) of positive TF regulators (correlation > 0.5, adjusted p-value < 0.01).

We then identified the positive TF regulators in Sertoli cells. Interestingly, the C1 cluster could be clearly distinguished from the other cell clusters as the 11 identified potential positive TF regulators either have highest or lowest activity in C1 (Figure 5D). For example, activity of several SP/KLF family members are enriched in C1, including SP1, which is reported to upregulate the transcription of nectin-2 and JAM-B in Sertoli cells (Lui and Cheng 2012). To validate this novel population, we searched for C1-specific markers. Notably, the C1 cluster showed a significantly higher gene activity of *Mbd3*, a 5hmC binding protein, when compared with other clusters (Figure 5E, Supplementary Figure S10A). Whole mount immunostaining of MBD3 protein confirmed the presence of this Sertoli cell subpopulation in mouse testis. The Sertoli cells could be indicated by Oct4-GFP- cells inside the seminiferous tubules, as all germ cells express Oct4-GFP at this stage. MBD3+ cells were Oct4-GFP- cells and exhibited patch-like distribution pattern deposits along the tubules, suggesting MBD3 only marked a subpopulation of Sertoli cells. Moreover, we only observed MBD3+ Sertoli cells in P1 seminiferous tubules, but not in P4 or P6 tubules (Figure 5F).

A recent study identified a potential Sertoli stem cell population in chicken testes characterized by expression of lower levels of *Dmrt1* and *Sox9 (Estermann et al. 2020)*. Consistent with this report, we found lower gene activity of Sertoli cell markers *Sox9*, *Dmrt1* and *Wt1* in the C1 (Figure 5E). In addition, FOS has the lowest motif binding activity in C1 (Supplementary Figure S10C), which is correlated with its inferred gene score (Figure 5E). Since FOS has been reported to mediate Sertoli cell differentiation, C1 cells might represent less differentiated Sertoli cells (Papadopoulos and Dym 1994).

### Characterization of TF regulation during perinatal Leydig cell development

Re-clustering of Leydig cells generated 3 main clusters (Figure 5G, Supplementary Figure S10D). Based on the gene activities of known Leydig cell markers, cluster C2 is likely to represent FLCs, as it expresses higher levels of *Gata4*, *Igf1* and *StAR*. C1 and C3 showed enrichment in different sets of SLC markers. C3, comprising mainly E18.5 and P0.5 cells, expressed higher levels of *Itgav* and *Nr2f2*, whereas C1, comprising mainly P5.5 cells, selectively expressed SLC markers *Pdgfra*, *Pdgfrb*, *Nes* and *Thy1* (Supplementary Figure S10E). 761 marker genes were identified across the clusters (FDR < 0.1, log2FC > 0.5), with most upregulated in C1 (Supplementary Figure S10F). GO terms associated with the marker genes in C1 include extracellular structure organization and connective tissue development (Supplementary Figure S10G). For instance, *Col4a1* and *Col4a2* showed higher gene activity in C1, accompanied by C1-specific peak-to-gene links to the bidirectional collagen IV promoter (Figure 5H). Previous studies have indicated that collagen IV-mediated signalling is involved in progenitor Leydig cell (PLC) proliferation, suggesting that C1 cells may represent a later developmental stage of SLC towards differentiating into PLCs (Anbalagan and Rao 2004).

Next, we identified the positive TF regulators of Leydig cells, which revealed 40 putative TFs (Figure 5I). For instance, *Gata4*, as a FLC marker, was consistently identified as a positive TF regulator upregulated in FLC cluster C2. *Ar*, which is crucial for the development of ALCs from SLCs, showed highest motif and gene activity in C1 (O’Shaughnessy et al. 2002). Thyroid hormone receptors *Thra* and *Thrb* were upregulated in C2. Other potential TF candidates include NR and KLF family members. These results suggested there is also large heterogeneity among different TFs in their involvement in different Leydig cell subpopulations.

Taken together, we uncovered potential genes and TFs that regulate the three main Leydig cell subpopulations present during the perinatal period.

### Stromal cell heterogeneity and PTM development during the perinatal period

Since PTMs and stromal cells were clustered into a single large cluster (Figure 1D) and PTMs are suggested to be derived from interstitial progenitors (Shen et al. 2020), PTMs and stromal cells were grouped together for subsequent analysis. Re-clustering of this cell group generated 10 clusters of cells (Supplementary Figure S11A and B). Within the clusters, C8, C9 and C10 showed higher gene activity of *Myh11*, indicating their PTM identity, while other clusters were stromal cells (Supplementary Figure S11C). 1927 marker genes were identified across the clusters (FDR < 0.1) (Supplementary Figure S11D). Notably, *Tcf21* is significantly upregulated in C2, while *Tcf21*+ cells were recently identified as bipotential somatic progenitors that could give rise to PTMs, Leydig cells and interstitial cells (Shen et al. 2020). Interestingly, the gene expression pattern of C2 also resembles telocytes, a recently described stromal cell type, which co-expresses *Cd34* and *Pdgfra* and is negative for *Pecam1*, *Kit* and *Acta2* (Supplementary Figure S11C) (Marini et al. 2018). Our results thus confirm that testis stromal cells are highly heterogeneous in nature.

Motif enrichment analysis of DARs across clusters indicated AR, PGR (progesterone receptor) and muscle-specific TF motifs were enriched in PTM clusters (Supplementary Figure S11E). In contrast, WT1 and SP family members exhibited higher TF motif activity in stromal cell cluster C3.

To identify the critical TFs involved in perinatal PTM development, we reconstructed the developmental trajectory and revealed 46 TFs with correlated gene scores and TF z-scores (Supplementary Figure S11F and G). Numerous candidates were associated with steroidogenesis (*Dlx6*, *Egr1*, *Xbp1*, *Sp3*) and spermatogenesis (*Hmga2*, *Nfya*, *Rfx1*, *Tbp*). Interestingly, 9 of the 46 TFs along the trajectory are homeobox genes. For example, *Hoxc6* is upregulated at the early stages and has been implicated in steroid hormone regulation (Ansari et al. 2011).

Lastly, we identified the putative positive TF regulators regulating PTM and stromal cells (Supplementary Figure S11H). Notably, *Smad3*, *Gata4*, *Tcf15*, *Nhlh2* and *Ppard* showed upregulation in PTM clusters, while *Rfx1*, *Rfx2*, *Lbx2*, *Nfya*, *Yy1* and *Fosl1* were upregulated in stromal clusters. Concordantly, *Gata4* has been described as a negative regulator of contractility in PTM, whereas *Smad3* is associated with androgen responsiveness and postnatal testis development (Itman et al. 2011; Wang et al. 2018).

### Identification of cell type-specific TFs in immune cells

Re-clustering of all the immune cells generated 4 cell clusters ((Figure 1D, Supplementary Figure S12A), which can be re-grouped as 3 main groups based on the expression of their corresponding marker genes, including T cells/NK cells (C1 and C2), myeloid cells (C3) and dendritic cells (C4) (Supplementary Figure S12B). Since testicular tissue-resident macrophages were shown to be involved in steroidogenesis, spermatogonia differentiation and Leydig cell function, our subsequent analysis focused on C3 which included the macrophage population. We first identified cluster-specific positive TF regulators (Supplementary Figure S12C and D). This accurately identified C/EBP proteins governing macrophage differentiation and mobilization and Kruppel-like factors (KLFs) controlling macrophage activation and polarization (McMahon et al. 1989; Mahabeleshwar et al. 2011; Date et al. 2014; Wada et al. 2015). Other well-known TFs with functions in myeloid cells/macrophages were identified including ZEB1, CREB1, FOSL2, EGR1 and NFAT5. Our analysis also revealed new candidates potentially regulating testis myeloid cells, such as NFYA, ZFP42 and TEF. Notably, NFYA and ZFP42 participate in testicular functions (Rezende et al. 2011; Iyer et al. 2016). TF candidates identified above could serve as future investigation targets related to testis immune cells.

### Single cell chromatin accessibility identified human GWAS target regulatory regions, genes and cell types in the testis

GWAS have been exceedingly successful in identifying nucleotide variations associated with specific diseases or traits. The significance of these findings can be realized only when the associated DNA sequence variation is linked to specific genes and the relevant cell types. Majority of the identified genetic variants are in the non-coding region of the genome, which include cell type specific enhancer regions. The potential role of these SNPs can be evaluated using the cell type specificity of these enhancers but such a map has not been generated for testis in the past. Therefore, we sought to predict which cell types in the testis may be the functional targets of polymorphisms from previous GWAS studies using the single cell level chromatin accessibility data and analytical framework as reported previously (Figure 6A) (Cusanovich et al. 2018). Cell type-specific LD score regression using testosterone level GWAS results revealed a significant increase in per-SNP heritability for testosterone level in the Leydig cell peak set (Figure 6B). Examining the variants overlapping Leydig cell DARs revealed rs4919685 (p=4.203e-15) and rs743572 (p=3.622e-07) located in the TSS-distal regions and the promoter of *Cyp17a1*, respectively (Figure 6C). CYP17a1 is one of the enzymes that converts testosterone to estradiol (Olson et al. 2007) and rs743572 has been suggested to be a functional SNP related to testosterone levels (Kakinuma et al. 2004). Similar analyses in male infertility and abnormal spermatozoa traits showed significant enrichment in SNP heritability in Sertoli cells (Figure 6B). In addition, we predicted immune cell types as the targets of GWAS variants related to several immune diseases, such as Lupus, Celiac disease, and Crohn’s disease. In contrast, the heritability of GWAS SNPs from traits not directly related to testis, such as Immunoglobulin A deficiency, Parkinson disease and major depressive disorder, were not enriched in any of the testicular cell types, indicating the robust nature of these results. Interestingly, we found that the GWAS variants related to coronary artery disease (CAD) were highly enriched in Sertoli cell and stromal cell peaks. This may be linked to the role of androgens in cardiovascular physiology as Sertoli cells are critical regulators of androgen secretion (Liu et al. 2003; Rebourcet et al. 2014).

**Figure 6.**
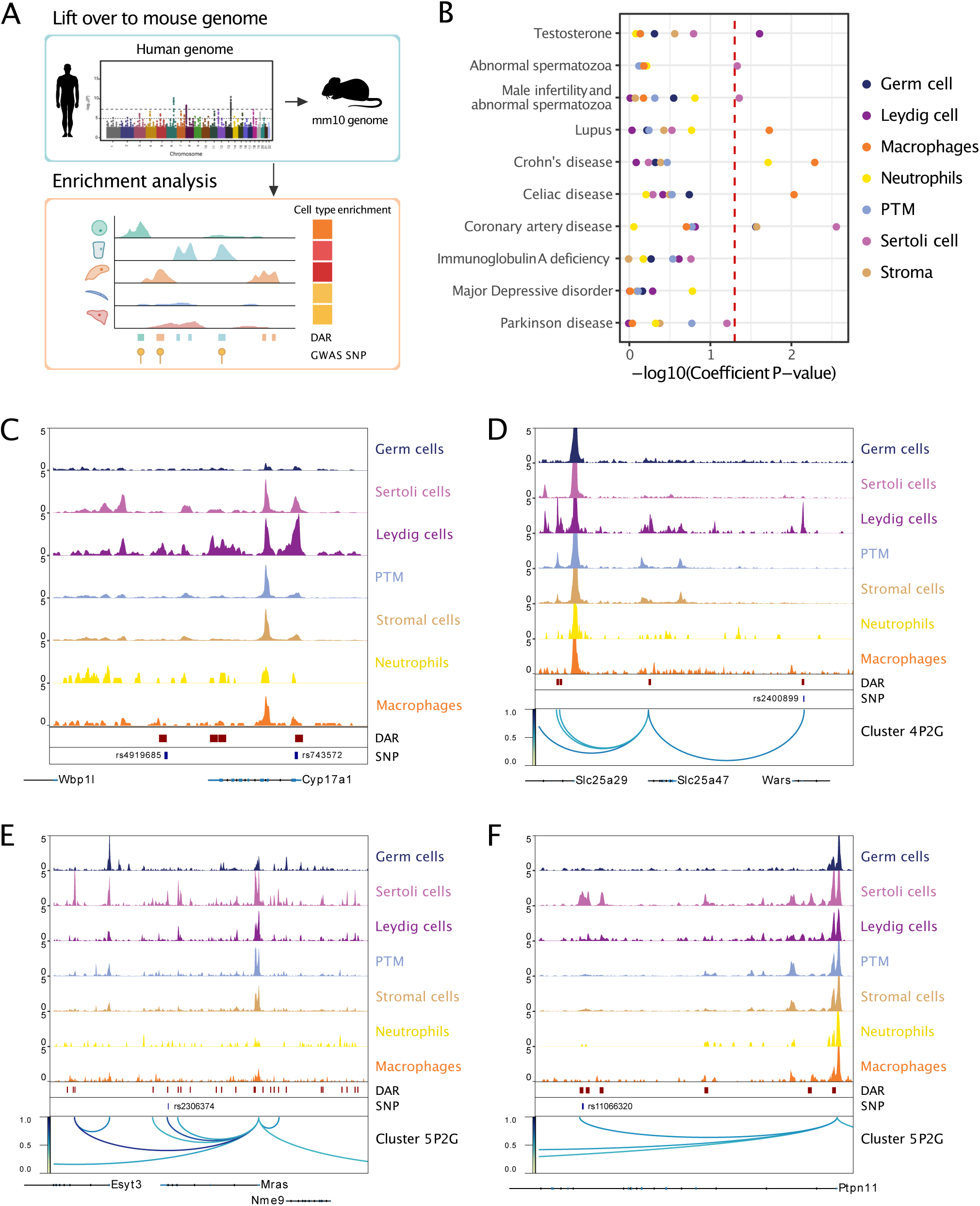
Single cell chromatin accessibility identified human GWAS targets in mouse testis. A. Illustration of heritability enrichment analysis. B. Enrichment of heritability for the selected traits within the cell type-specific DARs. C. Aggregated scATAC-seq profiles showing genetic variants (SNPs) overlapped with Leydig cell-specific DARs located at *Cyp17a1* locus. D. Aggregated scATAC-seq profiles showing SNPs overlapped with Leydig cell-specific DARs and peak-to-gene links to the *Slc25a47* locus. E. Aggregated scATAC-seq profiles showing SNPs overlapped with Sertoli cell-specific DARs and peak-to-gene links to the *Mras* locus. F. Aggregated scATAC-seq profiles showing SNPs overlapped with Sertoli cell-specific DARs and peak-to-gene links to the *Ptpn11* locus.

We also wanted to predict the genes that may be direct regulatory targets of a given non-coding polymorphism. We reasoned that their modulation of enhancer or promoter activity can be predicted by the co-accessibility framework. Using our catalog of cell type-specific peak-to-gene links, we linked TSS-distal GWAS variants to target genes. As an example of the Leydig cell-specific peak-to-gene link and testosterone level, we found the variant within *Wars* (rs2400899: p=4.993e-10) was connected to the nearby gene *Slc25a47*, which encodes a mitochondrial transporter (Figure 6D). This might be linked to the mitochondrial functions in regulating Leydig cell steroidogenesis (Hales et al. 2005). We then focused on Sertoli cells and CAD traits. We found the variant rs2306374 (p=6.49e-09) was located in a Sertoli cell-specific peak and was linked to *Mras*, which was reported to be associated with CAD (Figure 6E) (CARDIoGRAMplusC4D Consortium et al. 2013). We also found rs11066320 (p=1.03e-08) located in the *Ptpn11* locus (Figure 6F). The risk allele rs11066320 is robustly associated with HDL cholesterol levels, which are important in determining risk for CAD (Richardson et al. 2020).

Though not a focus of the current study, we note that the data generated here can also be used to help identify the gene targets of non-coding GWAS polymorphisms with a strong association with other traits not assayed above. For example, the genetic variant rs941576 lies within a conserved and regulatory region within intron 6 of the maternally expressed *Meg3* (Figure 3C). It has been suggested that rs941576 is strongly associated with T1D susceptibility and rheumatoid arthritis (Wallace et al. 2010; Wahba et al. 2020). Our data show that this SNP in the mouse genome lies within a peak linked to *Dlk1*. Therefore, it is plausible that this SNP alters the regulation of the paternally expressed *Dlk1*. Whether variation in *Dlk1* expression in testicular somatic cells can alter susceptibility to these diseased warrants further investigation. Taken together, a combination of our scATAC-Seq data with human traits or disease-relevant SNPs enables the prediction of target cell types and target genes for GWAS variants.

## Discussion

Understanding the genetic networks that underlie developmental processes requires a comprehensive understanding of the genes involved as well as the regulatory mechanisms that modulate the expression of these genes. To date there have only been a few studies adopting epigenetic approaches, including DNaseI-Seq and ChIP-Seq, to address the regulation of cell fate in somatic cells in the testis.

In this study, we present the first open chromatin map at single cell resolution for the developing testis. First, we showed that similar to scRNA-Seq, the chromatin accessibility information obtained from scATAC-Seq is able to define cell types in the testis. Second, identification of TFs has been challenging solely based on gene expression information from scRNA-Seq. Our scATAC-Seq provides additional information on TF regulation by correlating motif accessibility and predicted gene activity, which allows us to reveal cell type-specific TFs. Third, we found numerous peak-to-gene links among different cell types, which demonstrates the tight regulatory association between CREs and gene expression. More importantly, a significant amount of peak-to-gene links are within known testis enhancer regions, indicating the reliability of our analysis strategy. Lastly, our study illustrates the critical role of scATAC-Seq in identifying target cell types of known GWAS variants.

Our study revealed dynamic chromatin accessibility that tracks with germ cell differentiation. Notably, in the pseudotime trajectory analysis of germ cells, we observed two possible developmental trajectories from T1/T2-ProSGs to differentiating spermatogonia. Besides a conventional pathway from ProSG to differentiating spermatogonia through an undifferentiated spermatogonial state, we also observed a second trajectory in which T1-ProSG reached the differentiating state through two cell clusters with unknown identities. We speculated that this second trajectory may represent the germ cells undergoing the first wave of spermatogenesis, which are a subset of ProSGs that differentiate immediately into spermatogonia and continue to progress through spermatogenesis in early postnatal life (Yoshida et al. 2006). In fact, the low gene expression of *Ngn3* we observe along this trajectory is concordant with the observation that the first round of mouse spermatogenesis initiates directly from ProSGs without passing through the *Ngn3*-expressing undifferentiated spermatogonial stage. In contrast, the subsequent rounds of spermatogenesis are derived from *Ngn3*-positive undifferentiated spermatogonia (Yoshida et al. 2006), consistent with the high expression of *Ngn3* in the undifferentiated spermatogonia cluster in our dataset. Since there is currently a lack of marker genes to accurately identify gonocytes which undergo the first wave of spermatogenesis, our scATAC-Seq data uncovered a list of potential markers that warrants further investigation.

We also observed that scATAC-Seq was able to define novel cell populations not reported in published scRNA-Seq studies. For example, we identified a potential Sertoli stem cell population based on low expression of specific Sertoli cell markers. A recent study in chicken sex determination suggested a novel stem Sertoli population could be characterized by lower levels of *Dmrt1* and *Sox9*. We suggest that the MBD3+ cluster may represent a stem-cell-like subpopulation of Sertoli cells in mice that exists transiently shortly during the neonatal period (Estermann et al. 2020). This is in line with the observation that the 5hmC-binding protein MBD3 serves to maintain the pluripotency of ESCs (Yildirim et al. 2011; Hirasaki et al. 2018).

Cell type specific chromatin marks, when integrated with GWAS variants, can provide insights into disease causal cell types (Trynka et al. 2013). In the past, due to the scarce availability of such data in testis, it was difficult to assess the relevance of testicular cells in the pathobiology of various complex human traits. We showed that testicular cell type-specific peaks displayed increased heritability enrichment in cell populations consistent with the known biology and revealed new biological insights. For example, we observed that immune cells and Leydig cells demonstrate heritability enrichment for immune-related traits and testosterone levels, respectively. Intriguingly, Sertoli cells show higher heritability enrichment for CAD phenotypes, which leads us to hypothesize that where CAD-associated variants can act in Sertoli cells. The risk of CAD has been linked to testosterone level, increasing age and male gender (Nettleship et al. 2009). Our hypothesis can be supported by the role of Sertoli cells in testosterone level regulation through influencing the Leydig cell function and testicular vasculature (Rebourcet et al. 2016, 2017). Furthermore, co-accessibility measurement informs well-studied CAD-relevant genes, such as *Mras* and *Ptpn11* in Sertoli cells. These results define immediately testable hypotheses in which these variants modulate the activity of the cis-regulatory elements in Sertoli cells in the context of CAD. These findings thus suggested a new paradigm by which the Sertoli cells as a potential target of determinants of cardiovascular disease.

In summary, our high-resolution data has enabled detailed reconstruction of the gene regulatory landscape of testicular cell populations during key timepoints in testis development. Functional insight is further revealed by integrating multiple sources of genomic information and when combined provides an invaluable and unique resource for further investigation of key developmental events in the testis.

## Methods

### Animals

All the animal experiments were performed according to the protocols approved by the Animal Experiment Ethics Committee (AEEC) of The Chinese University of Hong Kong (CUHK) and followed the Animals (Control of Experiments) Ordinance (Cap. 340) licensed from the Department of Health, the Government of Hong Kong Special Administrative Region. All the mice were housed under a cycle of 12-hour light/dark and kept in ad libitum feeding and controlled the temperature of 22-24°C. Oct4-EGFP transgenic mice (B6; CBA-Tg(Pou5f1-EGFP)2Mnn/J, Stock no.: 004654) were acquired from The Jackson Laboratory (Ohbo et al. 2003). Oct4-EGFP transgenic mice and C57BL/6J mice were maintained in CUHK Laboratory Animal Services Centre.

### scATAC-Seq analysis

#### Sample collection

The testes of C57BL/6J mice at E18.5, P0.5, P2.5 and P5.5 were collected and with tunica albuginea removed. The testes were then digested with 1 mg/ml type 4 collagenase (Gibco), 1 mg/ml hyaluronidase (Sigma-Aldrich) and 5 µg/ml DNase I (Sigma-Aldrich) at 37°C for 20 min with occasional shaking. The suspension was passed through a 40-µm strainer cap (BD Falcon) to yield a uniform single cell suspension.

#### Cell lysis and tagmentation

Cell tagmentation was performed according to SureCell ATAC-Seq Library Prep Kit (17004620, Bio-Rad) User Guide (10000106678, Bio-Rad) and the protocol based on Omni-ATAC was followed (Corces et al. 2017). In brief, washed and pelleted cells were lysed with the Omni-ATAC lysis buffer containing 0.1% NP-40, 0.1% Tween-20, 0.01% digitonin, 10 mM NaCl, 3 mM MgCl2 and 10 mM Tris-HCl pH 7.4 for 3 min on ice. The lysis buffer was diluted with ATAC-Tween buffer that contains 0.1% Tween-20 as a detergent. Nuclei were counted and examined under microscope to ensure successful isolation. Same number of nuclei were subjected to tagmentation with equal ratio of cells/Tn5 transposase to minimize potential batch effect. Nuclei were resuspended in tagmentation mix, buffered with 1×PBS supplemented with 0.1% BSA and agitated on a ThermoMixer for 30 min at 37 °C. Tagmented nuclei were kept on ice before encapsulation.

#### scATAC-Seq library preparation and sequencing

Tagmented nuclei were loaded onto a ddSEQ Single-Cell Isolator (Bio-Rad). scATAC-Seq libraries were prepared using the SureCell ATAC-Seq Library Prep Kit (17004620, Bio-Rad) and SureCell ATAC-Seq Index Kit (12009360, Bio-Rad). Bead barcoding and sample indexing were performed with PCR amplification as follows: 37 °C for 30 min, 85 °C for 10 min, 72 °C for 5 min, 98 °C for 30 s, eight cycles of 98 °C for 10 s, 55 °C for 30 s and 72 °C for 60 s, and a single 72 °C extension for 5 min to finish. Emulsions were broken and products were cleaned up using Ampure XP beads. Barcoded amplicons were further amplified for 8 cycles. PCR products were purified using Ampure XP beads and quantified on an Agilent Bioanalyzer (G2939BA, Agilent) using the High-Sensitivity DNA kit (5067-4626, Agilent). Libraries were sequenced on HiSeq 2000 with 150 bp paired-end reads.

#### Sequencing reads preprocessing

Sequencing data were processed using the Bio-Rad ATAC-Seq Analysis Toolkit. This toolkit is a streamlined computational pipeline, including tools for FASTQ debarcoding, read trimming, alignment, bead filtration, bead deconvolution, cell filtration and peak calling. The reference index was built upon the mouse genome mm10. For generation of the fragments file, which contain the start and end genomic coordinates of all aligned sequenced fragments, sorted bam files were further process with “bap-frag” module of BAP (https://github.com/caleblareau/bap). Downstream analysis was performed with Harmony (Korsunsky et al. 2019) and ArchR (Granja et al. 2020). Fragment files were used to create the Arrow files in the ArchR package.

#### Clustering and gene score/transcription factor activity analysis

We filtered out low-quality nuclei with stringent selection criteria, including read depth per cell (>2,000) and TSS enrichment score (>20%). Potential doubles were further removed based on the ArchR method. Bin regions were cleaned by eliminating bins overlapping with ENCODE Blacklist regions, mitochondrial DNA as well as the top 5% of invariant features (house-keeping gene promoters). ArchR was used to estimate gene expression for genes and TF motif activity from single cell chromatin accessibility data. Gene scores were calculated using the addGeneScoreMatrix() function with gene score models implemented in ArchR. addDeviationsMatrix() function was used to compute enrichment of TF activity on a per-cell basis across all motif annotations based on chromVAR.

#### Trajectory analysis

Trajectory analysis was performed in ArchR. addTrajectory() function in ArchR was used to construct trajectory on cisTopic UMAP embedding. To perform integrative analyses for identification of positive TF regulators by integration of gene scores with motif accessibility across pseudo-time, we used the correlateTrajectories() function which takes two SummarizedExperiment objects retrieved from the getTrajectories() function.

#### Footprinting analysis

Differential transcription factor footprints across cell types were identified using the Regulatory Genomics Toolbox application HINT (Li et al. 2018). Aligned BAM files from different cell types were treated as pseudo-bulk ATAC-Seq profiles and then subjected to rgt-hint analysis. Based on MACS calling peaks, we used HINT-ATAC to predict footprints with the “rgt-hint footprinting” command. We then identified all binding sites of a particular TF overlapping with footprints by using its motif from JASPAR with “rgt-motifanalysis matching” command. Differential motif occupancy was identified with “rgt-hint differential” command and “–bc” was specified to use the bias-corrected signal.

### Statistical analysis

Assessment of statistical significance was performed using two-tailed unpaired t-tests, one-way ANOVA with Tukey multiple comparisons tests or Chi-squared tests. Statistical analysis was performed using GraphPad Prism v8. Associated P values are indicated as follows: *P < 0.05; **P< 0.01; ***P < 0.001; ****P < 0.0001; not significant (ns) P > 0.05.

## Data access

All raw and processed sequencing data generated in this study have been submitted to the NCBI Gene Expression Omnibus (GEO; https://www.ncbi.nlm.nih.gov/geo/) under accession number GSE164439.

## Acknowledgments

We thank Tommy Lo for scATAC-Seq experimental assistance and Jesse Xiao for interactive website development. We also thank the SBS Core Laboratories in The Chinese University of Hong Kong for technical support. This study was supported by General Research Fund (CUHK 14120418) to TLL.

## Author Contributions

JL, HC Suen, RMH and TLL conceived and designed the study. JL and HC Suen performed experiments, single-cell sequencing analysis and wrote the manuscript. SR, RZ and HC So performed LDSC analysis. JL, HC Suen, RMH and TLL analyzed and interpreted data. JL, HC Suen, ACSL, AWTL, THTC, MYC, HTC, RMH and TLL reviewed and edited the manuscript.

